# An antibody-tumor necrosis factor fusion protein that synergizes with oxaliplatin for treatment of colorectal cancer

**DOI:** 10.1101/698563

**Authors:** Davor Bajic, Kerry A. Chester, Dario Neri

## Abstract

We have cloned and characterized a novel fusion protein (Sm3E-TNF), consisting of the monoclonal antibody Sm3E in single-chain Fv fragment format, fused to murine tumor necrosis factor. The protein, which was expressed in mammalian cells and purified as a non-covalent stable homotrimer, bound to the cognate carcinoembryonic antigen (CEA) and retained tumor necrosis factor activity. A quantitative biodistribution experiment, performed in immunocompetent mice with CT26 colon carcinomas transfected with human CEA, revealed that Sm3E-TNF was able to preferentially accumulate in the tumors with excellent selectivity (tumor:blood ratio = 56:1, twenty-four hours after intravenous administration). The fusion protein mediated a rapid hemorrhagic necrosis of a large portion of the tumor mass, but a rim survived and eventually regrew. Surprisingly, the combination of Sm3E-TNF with 5-fluorouracil led to a reduction of therapeutic activity, while a combination with oxaliplatin led to a prolonged stabilization, with complete tumor eradication in 40% of treated mice. These therapy results were confirmed in a second immunocompetent mouse model of colorectal cancer (CEA-transfected C51 tumors) and provide a rationale for the possible clinical use of oxaliplatin in combination with fully-human antibody-TNF fusions.

## INTRODUCTION

Metastatic colorectal cancer remains an important unmet medical need as currently available treatment options are typically unable to induce long-term remissions(1,2). Immune checkpoint inhibitors remain inefficacious, with the exception of a small population of patients that have microsatellite instability (MSI) and deficient mismatch repair (dMMR)(3). These features are linked to an upregulation of PD-1 and other immune checkpoint molecules, providing rationale treatment with PD-1 blockers such as nivolumab and pembrolizumab in some cases, and suggesting that colorectal cancer may be amenable to other immunotherapeutic approaches (4).

Engineered cytokine products are gaining importance as a novel class of biopharmaceuticals for cancer immunotherapy(5–7), with a potential for combination with ionizing radiation(8–10), cytotoxic agents(11–16) and immune checkpoint inhibitors(17,18). In our experiments, we explore antibody-based delivery of tumor necrosis factor (TNF) to colorectal cancer, as an avenue to induce a rapid and selective tumor cell death within the neoplastic lesions while sparing normal organs. The approach has a sound platform; we and others have previously described TNF fusion proteins(19–22) and one of these products (L19-TNF) is currently being studied in pivotal clinical trials for the treatment of first-line metastatic soft-tissue sarcoma [NCT03420014; EudraCT Number: 2016-003239-38].

For a target we employed carcinoembryonic antigen (CEA), one of the best and most-validated tumor associated antigens for malignancies of epithelial origin(23). Importantly, a substantial body of nuclear medicine data has confirmed that metastatic colorectal cancer lesions can efficiently be reached *in vivo* by anti-CEA antibodies, both as intact immunoglobulins(24,25) and in antibody fragment formats(26,27). Sm3E, the targeting agent of choice, is an affinity-matured, humanized, anti-CEA single chain Fv antibody fragment (scFv), with an extremely low dissociation constant (Kd = 30 pM) and established tumor targeting potential in tumor-bearing mice (28). Antibody-TNF fusions specific to CEA have previously been described but exhibited suboptimal biodistribution and therapy results(22,29,30). We hypothesized that the use of a best-in-class antibody fragment and a mammalian cell expression system could improve *in vivo* performance, as *E.coli* production of antibody therapeutics has previously been associated with suboptimal tumor targeting results (31).

Here, we describe the generation and characterization of Sm3E-TNF, a mammalian-expressed fusion protein of Sm3E with murine TNF. We demonstrate selective targeting of Sm3E-TNF to CEA-transfected murine CT26 tumors in immunocompetent BALB/c mice models, leading to rapid hemorrhagic necrosis of the tumor mass. Moreover, our study showed that Sm3E-TNF exhibited potent anti-cancer activity as a single agent and was able to achieve complete tumor eradication in up to 80% of study animals when used in combination with oxaliplatin, an anti-cancer drug commonly used for the treatment of colorectal cancer(32).

## MATERIALS AND METHODS

### Cell lines

Chinese hamster ovary (CHO) cells, LM-fibroblasts, CT26 colon carcinoma cells and HT29 colorectal adenocarcinoma cells were obtained from the ATCC. The C51 colon carcinoma, which was kindly provided by M.P. Colombo, Istituto Nazionale Tumori, Milan, Italy). Cell lines were received between 2010 and 2017, expanded, and stored as cryopreserved aliquots in liquid nitrogen. Cells were grown according to the manufacturer’s recommendation. Authentication of the cell lines including check of post-freeze viability, growth properties and morphology, test for mycoplasma contamination, isoenzyme assay and sterility, were performed by the cell bank before shipment.

### Transfection of CEA into CT26 and C51 tumor cells

The gene for CEA was cloned into the mammalian expression vector pcDNA3.1(+) containing an antibiotic resistance for G418 Geneticin. CT26 wildtype cells were transfected with the pcDNA3.1(+) containing the human CEA gene using the TransIT-X2^®^ transfection reagent (Mirusbio) according to provider’s recommendations. Five days after the transfection, the medium was replaced with RPMI (10% Fetal Bovine Serum (FBS), 1% antibiotic-antimycotic (AA)) containing 0.5 mg/ml G418 (Merck) to select a stably transfected polyclonal cell line. A monoclonal cell line was generated from the stable polyclonal cell line by preparative single cell sorting performed using a BD FACSAria III. The process was tracked by FACS analysis. Different clones were expanded and further tested for antigen expression. Clone CT26 1D6 was selected for further use, on the basis of CEA expression which had been shown both by FACS and by immunofluorescence analysis. Similar procedures were used for the stable-transfection of C51 murine adenocarcinoma cells with human CEA, but electroporation with Amaxa™ 4D-Nucleofactor (Lonza) with the SG Cell Line 4D-Nucleofector® X Kit L (Lonza) was used for transfection. Relevant features and sequences of the vector used for cell transfection can be found in **Supplementary Figure 1**.

### FACS analysis

For cellular expression analysis of CEA, cells were detached with 50 mM EDTA and 5 × 10^5^ cells were stained with rabbit anti-CEA antibody (Sino Biological) (500 nM, 1 hour, 4°C) or Sm3E-TNF (500 nM, 1 hour, 4°C) in a volume of 100 μl FACS-Buffer (0.5% BSA, 2 mM EDTA in PBS). For signal amplification, the anti-CEA antibody was detected with an anti-rabbit AlexaFluor488 labeled antibody (Invitrogen) (1:200, 45 min, 4°C) whereas the Sm3E-TNF fusion protein was detected with a FITC labeled protein L (Acrobiosystems) (1:200, 45min, 4°C). In between and after staining, cells were washed with 100 μl FACS-Buffer and centrifuged at 300 rcf for 3 min. Stained cells were analyzed with a 2L-Cytoflex (Beckman-Coulter). When looking for CEA expression, HT29 cells were used as positive control while CT26 wildtype and C51 wildtype cells were used as negative control. FACS data were analyzed using the FlowJo 9 software suite (FlowJo LLC).

### Tumor models

CEA-transfected CT26 cells were grown to 80% confluence and detached with Trypsin-EDTA 0.05% (Life Technologies). Cells were then washed once with Hank’s Balanced Salt Solution (HBSS, pH 7.4, Gibco), counted and re-suspended in HBSS to a final concentration of 2.0 × 10^7^ cells/ml. Aliquots of 3 × 10^6^ cells, corresponding to 150 μl of the suspension, were injected subcutaneously in the right flank of female BALB/c mice (8-10 weeks of age, Janvier). Similar procedures were used for the CEA-transfected C51 cells but aliquots of 1 × 10^6^ cells in 100μl suspension were injected. Experiments were performed under a Project License granted by the Veterinäramt des Kantons Zürich, Switzerland (27/2015).

### Cloning, expression, and *in vitro* characterization of the fusion protein Sm3E-TNF

The VL and VH genes (US 2008/0003646 A1) of an affinity matured humanized anti-CEA monoclonal antibody (Sm3E)(33) were fused through a G_4_S-linker sequence. The resulting VH-(G_4_S)_3_-VL ScFv fragment was further fused at the N-terminus of the murine TNF gene through a S_4_G-linker and the final construct VH-(G_4_S)_3_-VL-(S_4_G)_3_-TNF was then cloned into the mammalian expression vector pcDNA3.1 (+) vector (Invitrogen). A VH-(G_4_S)_3_-VL ScFv fragment specific for hen egg lysozyme (KSF)(34), used as negative control, was fused at the N-terminus of the TNF as described for the Sm3E construct. The sequences for both fusion proteins can be found in **Supplementary Figure 2 and 3**.

The fusion proteins were expressed in CHO cells by transient gene expression, using previously-described procedures(35), and were purified to homogeneity using either protein A (KSF-TNF; Sino Biological) or protein L (Sm3E-TNF; Thermofisher) chromatography. The purified proteins were characterized by SDS-PAGE (Invitrogen), size-exclusion chromatography (Superdex200 10/300GL, GE Healthcare) and protein mass spectrometry (UPLC-ESI-ToF-MS, Waters). The biological activity of Sm3E-TNF was assessed by incubation with mouse LM fibroblasts or CEA-transfected CT26 cells. In this assay, cells were incubated in 96-well titer plates containing medium supplemented with 2μg/ml Actinomycin D and varying concentration of fusion protein. After 24h at 37°C, cell viability was determined using a Cell Titer Aqueous One Solution kit (Promega).

### Biodistribution studies

The *in vivo* tumor targeting performance of the fusion proteins was assessed by quantitative biodistribution after radiolabeling following previously published experimental procedures(36). Briefly, approximately 7μg of radioiodinated Sm3E-TNF or KSF-TNF were injected into the lateral tail vein of CEA-transfected CT26 tumor-bearing BALB/c (Janvier) mice (n = 5). Animals were sacrificed by CO_2_ asphyxiation 24 hours after injection. Organs and tumors were excised, weighed and the corresponding radioactivity was measured using a Cobra γ counter (Packard). Biodistribution results were expressed as percent of injected dose per gram of tissue (%ID/g ± Standard Error of the Mean).

### H&E staining and Immunofluorescence studies

Mice bearing CEA-transfected CT26 tumors were injected with a single dose of fusion protein alone or in combination with oxaliplatin according to the therapy schedule and subsequently sacrificed at variable time points after injection for microscopic analysis. Tumors were excised and embedded in cryoembedding medium (Thermo Scientific). Ice-cold acetone-fixed cryostat sections (10μm) were prepared and stained with hematoxylin and eosin(H&E) (Sigma-Aldrich) using routine methods, or subjected to immunofluorescence analysis. For immunofluorescence, the acetone-fixed sections were stained using the following antibodies: rat anti-CD31 (BD Bioscience; 553370), goat anti-CD31 (R&D Systems; AF3628), rabbit anti-activated-caspase-3 (Sigma; C8487), rat anti-NKp46 (Biolegend; 137601), rat anti-CD4 (Biolegend; 100423), rat anti-CD8 (Biolegend; 100702), rat anti-FoxP3 (eBioscience; 14-5773-82), and detected with anti-goat AlexaFluor594 (Invitrogen; A11058), anti-rat AlexaFluor594 (Invitrogen; A21209), anti-rat AlexaFluor488 (Invitrogen; A21208), anti-rabbit AlexaFluor488 (Invitrogen; A11008). Cell nuclei were counterstained with DAPI (Invitrogen; D1306). Slides were mounted with fluorescent mounting medium and analyzed with Axioskop2 mot plus microscope (Zeiss)

### Therapy studies

Tumor growth was monitored daily and a caliper was used to determine the tumor volume (volume= 0.5 × length × width^2^). When tumors had reached a volume of approximately 100mm^3^, mice were randomly subdivided into distinct study groups (n=5). Sm3E-TNF (4 μg), dissolved in phosphate-buffered saline (PBS), was administered every 48 hours by injection into the lateral tail vein. Vehicle control animals received the same volume of PBS alone. Chemotherapeutic agents [5-Fluorouracil (50mg/kg; Sandoz Pharmaceuticals AG) or oxaliplatin (7.5mg/kg; Sandoz Pharmaceuticals AG)], dissolved in PBS, were injected into the lateral tail vein only once, 30 minutes prior to the first fusion protein administration.

### AH1-Tetramers analysis

The presence of AH1-specific T-cells in draining lymph nodes (LN) and tumors of mice treated with saline, oxaliplatin and Sm3E-TNF alone or in combination with oxaliplatin was assessed by flow cytometry analysis, following previously published experimental procedures(37)

### Statistical analysis

Data were analyzed using Prism 7.0 (GraphPad Software, Inc.). Statistical significance was determined with a regular 2-way ANOVA test with a Bonferroni posttest. Data represent mean± SEM, P < 0.05 was considered statistically significant (*, P < 0.05; **, P<0.01; ***, P≤0.001; and ****, P < 0.0001).

## RESULTS

### Expression, characterization and in vitro analysis of TNF fusion proteins

We cloned and expressed Sm3E-TNF in Chinese hamster ovary (CHO) cells. The fusion protein featured the antibody in scFv format(38), leading to the formation of a stable non-covalent homotrimeric structure, mediated by the murine TNF moiety [**Figure 1A**]. The product was purified to homogeneity using Protein L chromatography, as assessed by SDS-PAGE analysis, size-exclusion chromatography and mass spectrometry [**Figure 1B-D**]. In parallel, we produced KSF-TNF, a negative control fusion protein comprising an scFv specific to hen egg lysozyme fused to murine TNF [**Supplementary Figure 4**]. Sm3E-TNF was able to kill TNF-sensitive LM1 fibroblasts (as well as CEA-transfected CT26 cells) *in vitro* at picomolar concentrations [**Figure 1E**]. Stably-transfected murine CT26 and C51 colorectal cancer cell lines, expressing human CEA (the cognate antigen recognized by Sm3E) had been produced by conventional methodologies, as described in the *Materials and Methods* section. Specific binding of Sm3E-TNF (but not KSF-TNF) to these cell lines was confirmed by FACS analysis, using fluorescently-labeled protein L as detection reagent [**Figure 2**].

**Figure 1:**
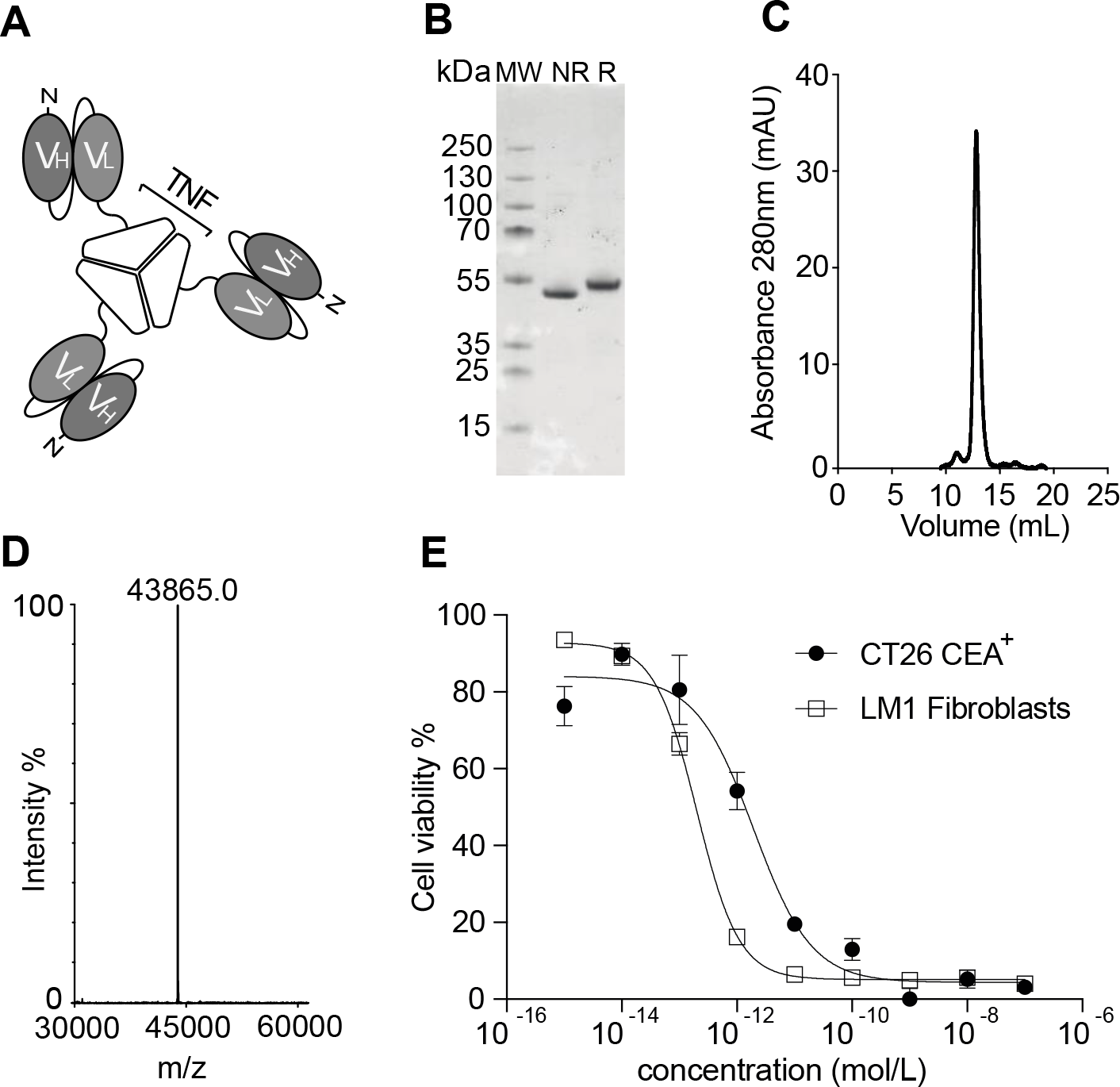
Cloning, expression and characterization of Sm3E-TNF and its Tumor-targeting properties. A) Schematic representation of Sm3E-TNF in noncovalent homotrimeric format. B) SDS-PAGE analysis of purified Sm3E. MW, molecular weight; R, reducing conditions; NR, non reducing conditions. C) Size exclusion chromatography profile. D) ESI-MS profile. F) TNF bioactivity assay, based on the killing of LM1 fibroblasts and of CEA transfected CT26 cells respectively.

**Figure 2:**
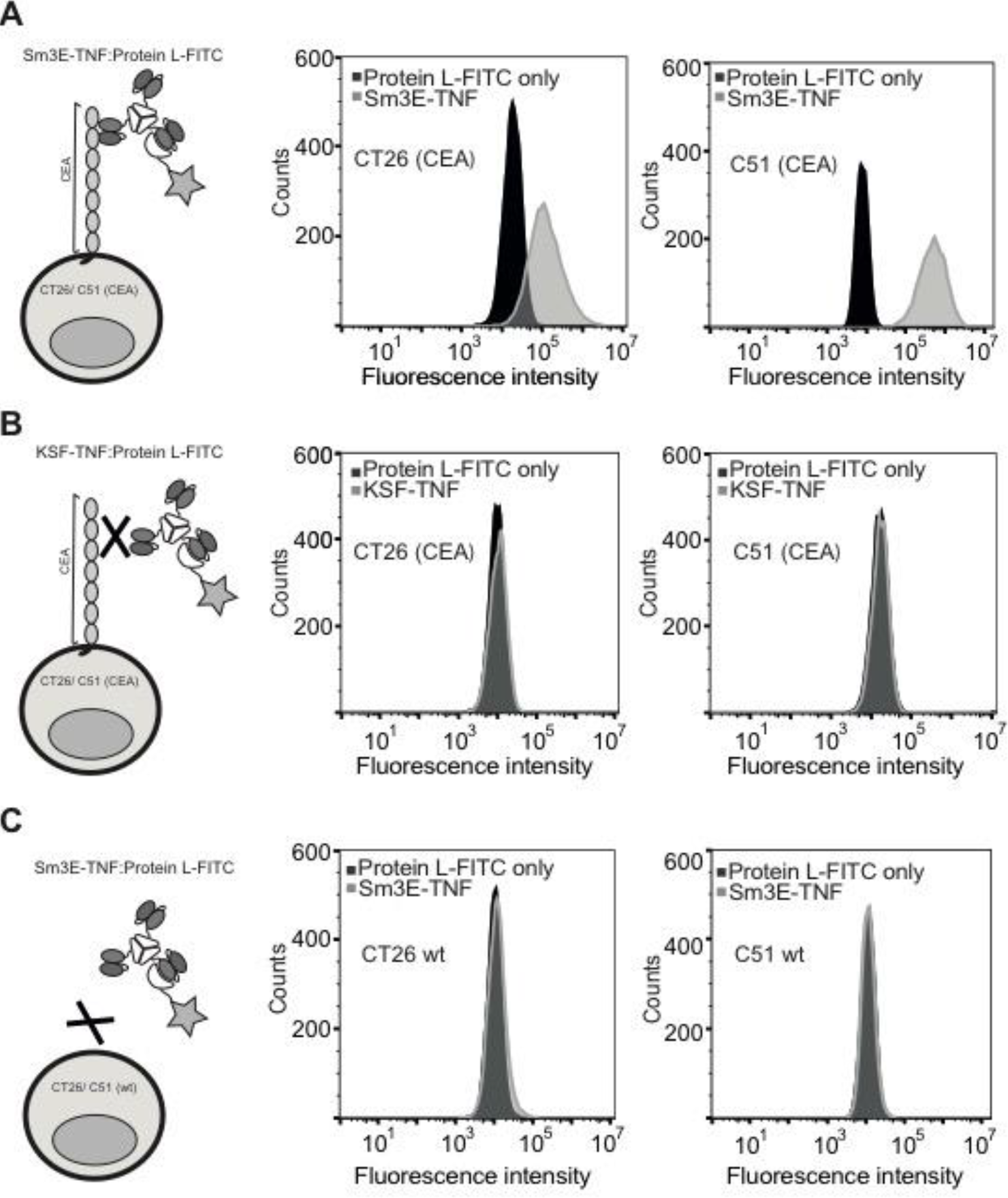
Flow cytometry analysis of Sm3E-TNF and KSF-TNF binding on CEA-transfected CT26, CEA-transfected C51, CT26 wildtype and C51 wildtype cells. A) Sm3E-TNF was bound to the CEA-transfected CT26 and CEA-transfected C51 cells and the signal was amplified with a FITC-labeled Protein L, whereas the negative control KSF-TNF (B) was not binding to the CEA-transfected CT26 and CEA-transfected C51 cells. C) Sm3E-TNF was not binding to the CT26 and C51 wildtype cells showing that the fusion-protein was specific to CEA.

### In vivo localization to CEA- expressing tumors and monotherapy with Sm3E-TNF

Twenty-four hours after intravenous administration to mice, Sm3E-TNF preferentially localized to CEA-expressing CT26 murine colorectal tumors *in vivo*, as assessed by a quantitative biodistribution analysis with radiolabeled protein preparations where Sm3E-TNF had a 3.52 ± 0.38 % inj. dose/g and tumor to blood ratio of 55.85 ± 10.42, whereas KSF-TNF had a 0.74 ± 0.28 % inj. dose/g and tumor to blood ratio of 3.46 ± 1.34 [**Figure 3**].

**Figure 3:**
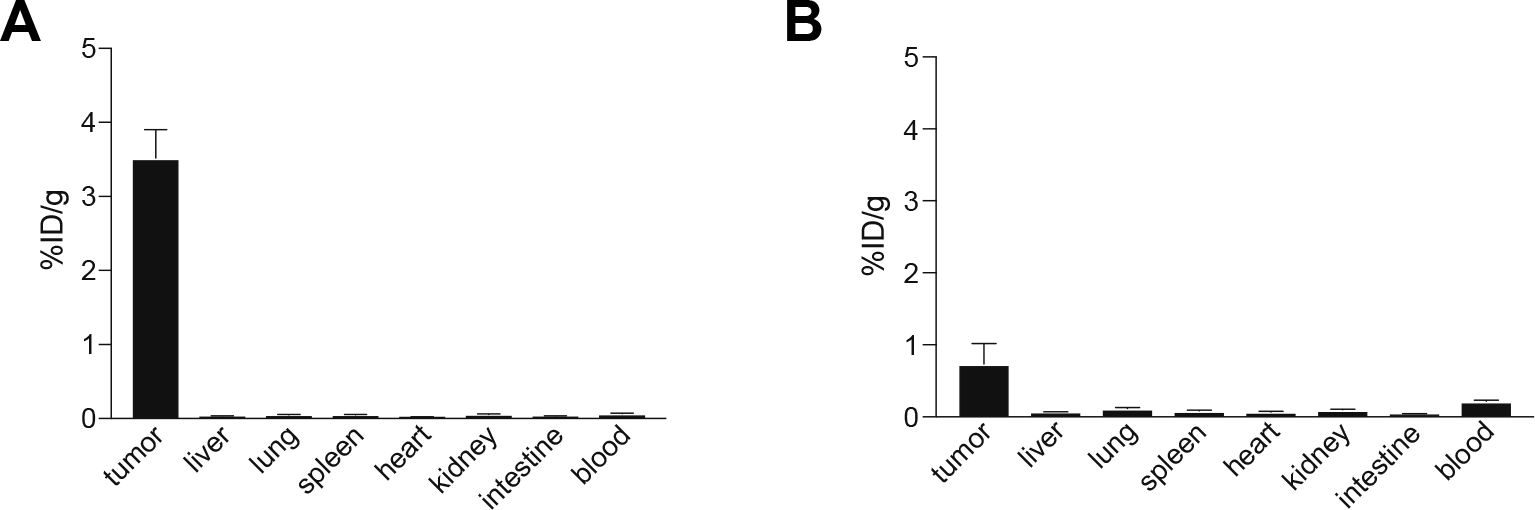
Quantitative biodistribution analysis of radioiodinated Sm3E-TNF (left) and KSF-TNF(right) in immunocompetent Balb/c mice bearing CEA-transfected CT26 coloncarcinoma tumors. Approximately seven μg of radioiodinated fusion protein was injected intravenously into the lateral tail vein and mice were sacrificed 24 hours after injection, organs were excised, weighted and the radioactivity of organs and tumors was measured. Results are expressed as percentage of injected dose per gram of tissue (%ID/g ± SEM; n= 5mice per group).

A first set of monotherapy experiments confirmed that the tumor-homing Sm3E-TNF product exhibited a more potent anti-cancer activity than KSF-TNF, with a significant difference (P≤0.001) observed for slow growing CEA-expressing CT26 tumors. However, at the dose used (three injections of 4 μg of Sm3E-TNF), cancer could not be cured in a monotherapy regimen, which was associated with a body weight loss of ~5% [**Figure 4**].

**Figure 4:**
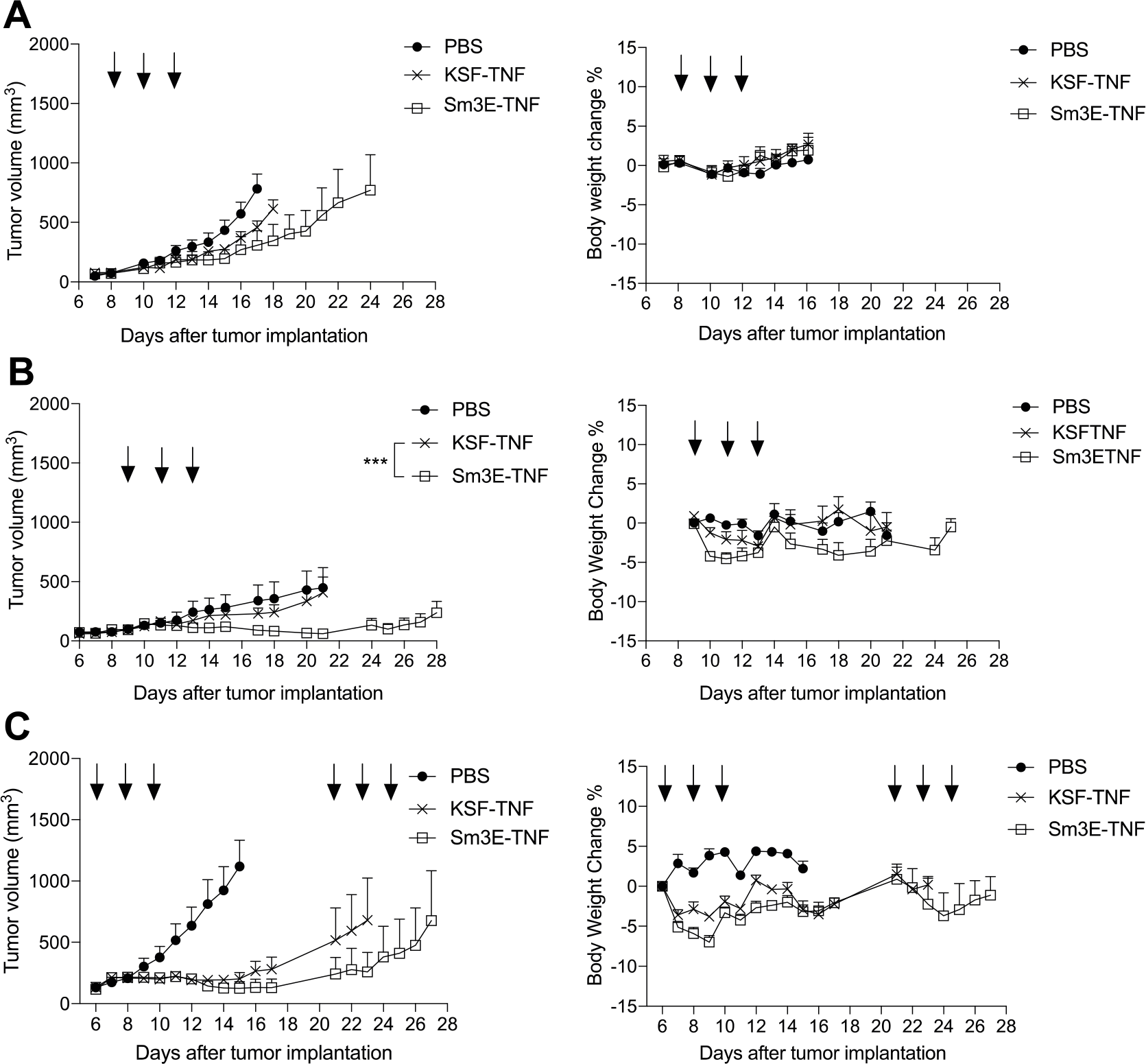
Comparison of therapeutic performance and tolerability, expressed as Body Weight Change (left panels), of single agent Sm3E-TNF versus KSF-TNF for the treatment of colorectal cancer, in balb/c mice bearing CEA transfected CT26 or C51 colorectal tumors. Sm3E-TNF exhibited a more potent anti-cancer activity compared to KSF-TNF, treatment started when tumors reached a volume of 100 mm^3^ and mice were randomly grouped. Data represent mean tumor volume ± SEM, n = 5 mice per group; CR, complete response. P < 0.05 was considered statistically significant (*, P < 0.05 and ***, P≤0.001). **A**, Fast growing CEA-transfected CT26 clone, three injections, one every 48 hours, of either PBS, 2μg Sm3E-TNF or 2μg KSF-TNF. **B**, Slow growing CEA-transfected CT26 clone and **C**, CEA-transfected C51 tumors, three injections, one every 48 hours, of either PBS, 4μg Sm3E-TNF or 4μg KSF-TNF., three injections of either PBS, 4μg Sm3E-TNF or 4μg KSF-TNF (injections are indicated by arrows).

### Combination studies

Chemotherapeutic strategies for the treatment of metastatic colorectal cancer in patients typically make use of 5-fluorouracil (5-FU) and/or oxaliplatin(32). For this reason, we performed a first set of experiments in CEA-expressing CT26 cells, using these drugs alone or in combination with Sm3E-TNF. Surprisingly, the combination of 5-FU (50 mg/Kg) with Sm3E-TNF resulted in a worse therapeutic performance than single agent Sm3E-TNF [P value < 0.0001] [**Figure 5A**]. By contrast, the combination with oxaliplatin (7.5 mg/Kg) strongly potentiated the action of Sm3E-TNF, leading to cancer cures in 40% of the animals [**Figure 5B**]. Cured mice rejected a subsequent challenge with tumor cells at day 50, indicating the onset of a protective immunity (data not shown). Similar therapy experiments were also performed in CEA-transfected C51 tumors, with the aim to investigate whether Sm3E-TNF (alone or in combination with oxaliplatin) would be active in this model. The fusion protein was able to potently inhibit tumor growth when used as monotherapy. Importantly, when Sm3E was combined with oxaliplatin, 80% of tumor-bearing mice were cured [**Figure 5C**].

**Figure 5:**
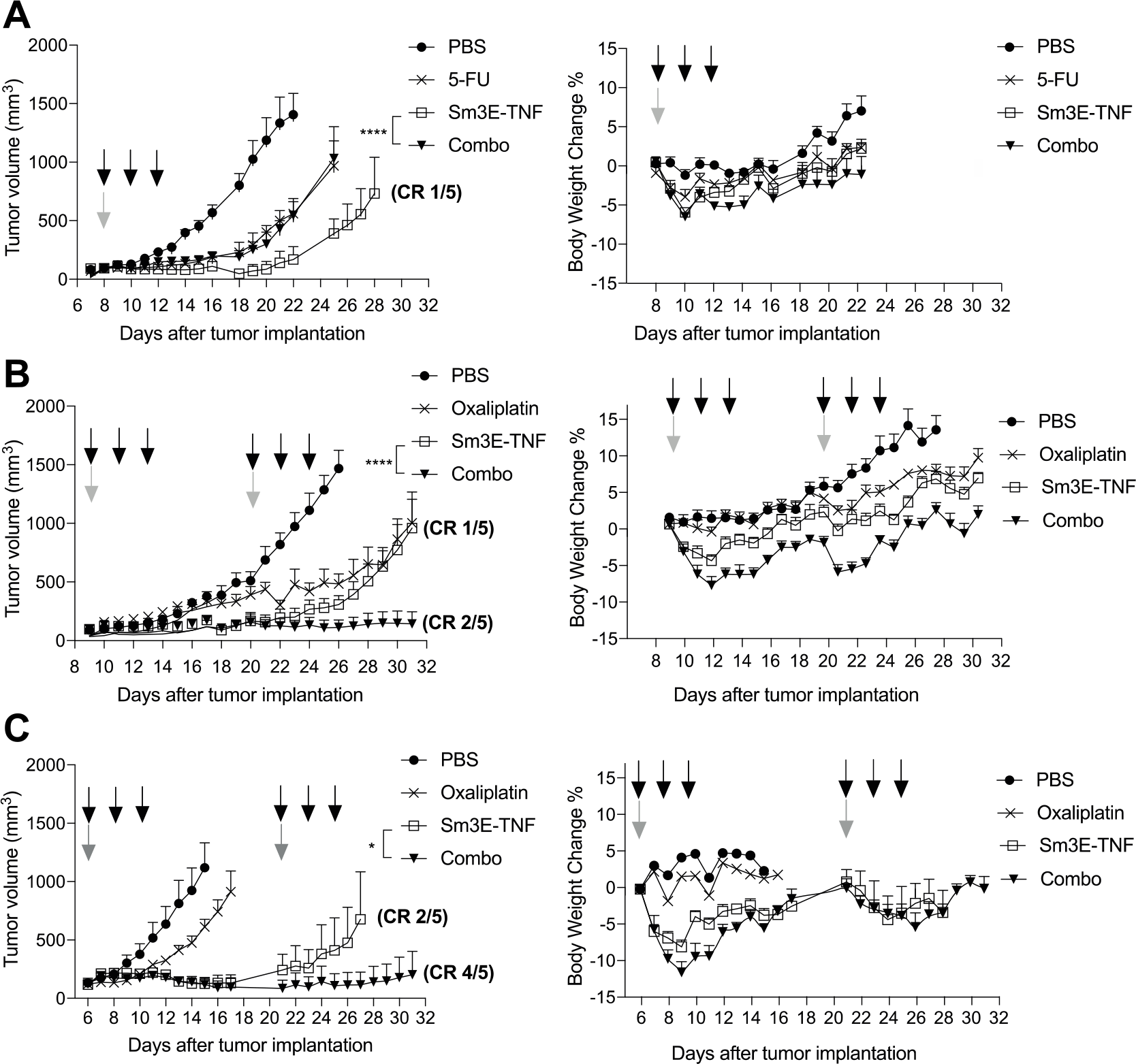
Therapeutic performance and tolerability, expressed as Body Weight Change (left panels), of Sm3E-TNF alone or in combination with standard chemotherapeutics for the treatment of colorectal cancer, in balb/c mice bearing CEA transfected CT26 or CEA transfected C51 coloncarcinoma tumors. Data represent mean tumor volume ± SEM, n = 5 mice per group; CR, complete response. P < 0.05 was considered statistically significant (*, P < 0.05 and ****, P < 0.0001). A) Comparison of single agents Sm3E-TNF, 5-Fluorouracil or combination of 5-Fluorouracil and Sm3E-TNF against CEA transfected CT26 tumors. The treatment started when tumors reached a volume of 100 mm^3^, mice were randomly grouped and injected in the tail lateral vein with a single injection of 50mg/kg 5-Fluorouracil, three injections, one every 48 hours, of 4μg of Sm3E-TNF, or the combination where the 5-Fluorouracil was administered 30 minutes prior of the fusion protein (injections are indicated by arrows. Grey, 5-Fluorouracil; black, Sm3E-TNF). PBS was used as vehicle. Comparison of single agents Sm3E-TNF, oxaliplatin or the combination against CEA transfected CT26 tumors (B) and CEA transfected C51 tumors (C), treatment started when tumors reached a volume of 100 mm^3^, mice were randomly grouped and injected in the tail lateral vein with a single injection of 7.5mg/kg oxaliplatin, three injections, one every 48 hours, of 4μg of Sm3E-TNF, or the combination where the oxaliplatin was administered 30 minutes prior of the fusion protein. A second cycle of injections was performed once the mice recovered the weight lost during the first cycle (injections are indicated by arrows. Grey, oxaliplatin; black, Sm3E-TNF). PBS was used as vehicle.

### Preliminary mechanistic evaluation

From a mechanistic viewpoint, the antibody-based delivery of TNF caused a rapid and selective hemorrhagic necrosis in both CEA-transfected CT26 and C51 tumors [**Figure 6A**], similar to that previously described by our group using different TNF fusion proteins directed against components of the tumor extracellular matrix (11,19). A microscopic analysis of H&E-stained sections revealed that the tumor architecture was intact after treatment with saline or oxaliplatin, but was largely necrotic in both Sm3E-TNF monotherapy groups and in the combination groups [**Figure 6B**]. The induction of hemorrhagic necrosis was less efficient when KSF-TNF was used, leaving groups of living cells within the tumor mass [**supplementary 5**].

**Figure 6:**
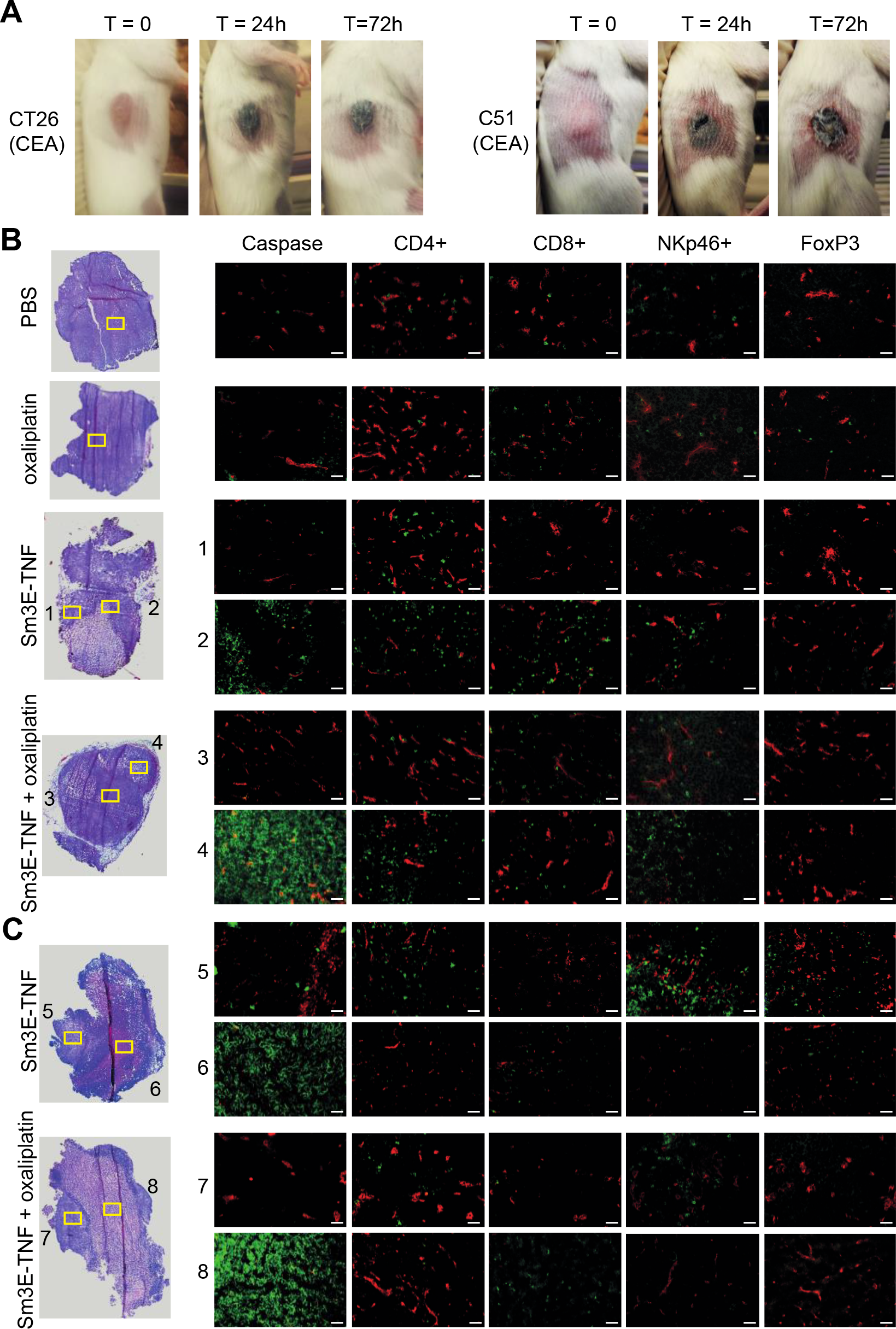
Sm3E-TNF-induced Hemorrhagic necrosis in tumors. A) Representative images of a targeted-TNF treated tumor-bearing mice before, 24hours and 72 hours after intraveneous injection of Sm3E-TNF (left CEA-transfected CT26; right CEA-transfected C51). The black color of the tumor is a typical macroscopic effect of hemorrhagic necrosis. B&C) From left: Hematoxylin & eosin analysis and detection of apoptosis (activate caspase 3), CD4^+^ T cells (CD4), CD8^+^ T cells (CD8), NK cells (NKp46), and Tregs (FoxP3) of sequential sections of CEA-transfected CT26 tumors by immunofluorescence (green, Alexa Fluor 488). Blood vessels stained with anti-murine CD31 (red, Alexa Fluor 594). Mice have been injected once with either PBS, oxaliplatin, Sm3E-TNF alone or in combination with oxaliplatin and sacrificed after 24h (B) or 72h (C). H&E original magnification x5; immunofluorescence original magnification x20. Scale bar = 100μm. The yellow rectangles on the H&E tumor sections define the approximate area of the sequential sections analyzed by immunofluorescence.

We also performed a time series of microscopic investigations of CEA-transfected CT26 tumors following treatment, in order to assess structural variations by H&E staining and by immunofluorescence analysis [**Figure 6**]. The tumor mass was almost completely necrotic 24 hours after Sm3E-TNF therapy. Serial tissue sections were studied by immunofluorescence and areas corresponding to rectangles in the H&E sections. The most striking differences between saline, oxaliplatin and Sm3E-TNF (monotherapy or combination) treatment regimens related to the ability of targeted TNF to induce hemorrhagic necrosis and to promote an increased intratumoral density of NK cells and of T cells.

The AH1 retroviral antigen is frequently the most relevant rejection antigen in BALB/c derived tumors(11,37) and we hypothesized that AH1 might play a role in the tumor response. However, although we detected AH1-specific T cells in the CEA-transfected tumors using tetramer reagents [**Supplementary Figure 6**], there was no apparent link to a treatment regime.

## DISCUSSION

Sm3E-TNF is a novel fusion protein, capable of specific recognition of CEA and of selective homing to CEA-expressing colorectal tumors *in vivo*. The product was able to induce a potent tumor growth retardation, which was enhanced by combination with oxaliplatin. Mice which were cured as a result of the combination treatment rejected subsequent challenges of the same tumor cells at day 50, indicating the onset of protective anticancer immunity.

From a pharmacodynamic viewpoint, the most visible effect of Sm3E-TNF on CEA-positive tumors was the rapid induction of hemorrhagic necrosis, leading to the death of large tumor portions after a single injection of the product. Living cells, however, typically survived at the tumor rim, with an enhanced but patchy infiltration by T cells and NK cells.

We have previously shown that the targeted delivery of TNF to colorectal tumors and to other cancer types in immunocompetent mouse models potently synergizes with other therapeutic modalities, including other cytokines (e.g., IL2(43) and IL12(44)), cancer vaccines(37) and immune checkpoint inhibitors(45). In the current manuscript, we studied the combination Sm3E-TNF with 5-FU and with oxaliplatin, as these drugs are commonly used in chemotherapeutic strategies for the treatment of patients with metastatic colorectal cancer(46). Surprisingly, 5-FU worsened the therapeutic effect of Sm3E-TNF, while the combination with oxaliplatin improved anti-cancer activity. Oxaliplatin has been shown not only to induce direct tumor cell death by apoptosis and secondary necrosis, but also to potentiate anti-cancer immunity through immunogenic cell death (47–49). In a previous vaccination experiment, CT26 cells were treated *in vitro* with oxaliplatin for 24 hours and inoculated subcutaneously in immunocompetent BALB/c mice, which upon re-challenge rejected a new subcutaneous injection of living CT26 tumor cells. In a further experiment, it was shown that immunocompetent tumor-bearing mice treated with oxaliplatin responded better to the treatment than immunocompromised tumor-bearing mice, demonstrating that the anticancer effects were, at least in part, mediated by the immune system(47,50). CT26 and C51 tumors have previously been described as rather immunogenic tumors, by virtue of a higher proportion of infiltrating T cells and NK cells (51,52). Tumors of BALB/c origin exhibit an unusually high infiltration by CD8+ T cells which recognize the AH1 peptide, derived from the aberrantly-expressed envelope protein of murine leukemia virus(37,53). Recent studies have indicated that non-coding regions (including retroviral antigen sequences) may represent the main source of tumor rejection antigens(54).

Interestingly CEA-expressing cell lines are not rejected by the mouse immune system after subcutaneous injection. This immunological tolerance has also been observed in previous studies, where antigens of human origin were transfected in murine cell lines that have been subsequently used for tumor implantation in syngeneic immunocompetent mice(39–41). In one of these studies, the *ex vivo* analysis of CEA-transfected tumor bearing mice showed the presence of anti-CEA antibody in the blood, however the tumors were still able to grow and the role of these antibodies in the inhibition of the tumor growth was not identified(41). One of the reasons for which the murine immune system fails to clear a potentially immunogenic human antigen-transfected cell line might be due to the tumor immunosuppressive microenvironment. This unfavorable microenvironment could prevent the host immune system from rejecting the tumor, for instance, by tumor-induced impairment of antigen presentation, where the immunosuppressive microenvironment causes the antigen presenting cells to present tumor antigens that lead to the induction of T-cell tolerance(42).

Various immunological approaches have been attempted for the treatment of metastatic colorectal cancer, but unfortunately clinical success has been limited. A bispecific antibody, directed against CEA and CD3, showed promising results in preclinical models, but did not mediate objective responses in patients with colorectal cancer(55). Similarly, anti-CEA antibodies have been used to deliver a human IL2 mutant to colon tumors(56). The targeted delivery of TNF to CEA-positive lesions may represent an alternative and potentially more efficacious strategy, because of the rapid induction of tumor cell death.

The debulking of the tumor mass triggered by Sm3E-TNF in mouse models is impressive, but it is still unclear whether similar effects could be observed in patients. Isolated limb perfusion (ILP) of patients with an advanced melanoma and soft-tissue sarcoma of the limbs, who were candidates for amputation, using TNF in combination with melphalan and mild hyperthermia, typically induces an hemorrhagic necrosis of the neoplastic mass(57–59), but TNF is administered at doses which would not be compatible with systemic administration. The targeted delivery of TNF to the tumor site, mediated by fusion with the L19 antibody (specific to the alternatively-spliced EDB domain of fibronectin) has recently been shown to induce complete responses and rapid hemorrhagic necrosis in the ILP setting, at doses which are more than 10-times lower than those conventionally used with recombinant human TNF(59). There is an urgent need to perform similar studies in patients with disseminated disease, following intravenous administration of antibody-TNF fusions. Perfusion MRI(60) or ^18^F-FDG-PET(61) procedures may be considered as imaging modalities to visualize vascular shutdown of the tumor, immediately after the administration of targeted TNF therapeutics.

In the pre-clinical studies presented, the humanized Sm3E antibody was fused to murine TNF because the human cytokine homologue is only partially reactive with murine TNF receptors(62). It should be straightforward, however, to fuse Sm3E with human TNF, in full analogy to that previously described for L19-TNF(59,63). A human Sm3E-TNF fusion protein might be expected to effectively deliver TNF to target, as CEA-specific antibodies have been extensively validated for their ability to selectively localize to metastatic colorectal cancer lesions using Nuclear Medicine procedures(26,27,64,65). Our results indicate that a fully human Sm3E-TNF product could have potential to help debulk cancer lesions, and facilitate the management of patients with metastatic disease, in combination with oxaliplatin.

## ACKNOWLEDGEMENTS

We gratefully acknowledge financial support from the ETH Zürich, the Swiss National Science Foundation (grant number 310030B_163479/1) and the European Research Council (ERC, under the European Union’s Horizon 2020 research and innovation program, grant agreement 670603). We would like to thank Dr. Roberto De Luca (Philochem AG, Otelfingen) and Dr. Mattia Matasci (Philochem AG, Otelfingen) for their help with experimental procedures.

